# Hazard detection with monocular bioptic telescopes in a driving simulator

**DOI:** 10.1101/355925

**Authors:** Xiaolan Tang, P. Matthew Bronstad, Lauren Spano, Amy L. Doherty, Mojtaba Moharrer, Eli Peli, Alex R. Bowers

**Affiliations:** Schepens Eye Research Institute, Massachusetts Eye and Ear, Harvard Medical School, Boston MA; College of Information Engineering, Capital Normal University, Beijing 100048, China

**Author notes:** Corresponding author: Alex Bowers.

**Keywords:** Bioptic telescope, scotoma, hazard detection, low vision, driving, central vision loss

## Abstract

ABSTRACT

**Purpose:** Recently we developed a driving simulator paradigm to evaluate detection of road hazards when using a bioptic telescope and conducted an initial study using normally-sighted observers with simulated vision loss. We now extend our investigation to quantifying the extent to which visually impaired bioptic users are able to use their fellow (non-telescope) eye to compensate for the ring scotoma of a monocular bioptic telescope. We tested the hypothesis that detection rates would be higher in binocular viewing (fellow eye could potentially compensate) than monocular viewing (fellow eye patched so it could not compensate) for pedestrian hazards present in the scene only while the telescope was being used.

**Methods:** Sixteen bioptic telescope users (17-80 y) completed six test drives, including three with binocular viewing interleaved between three with monocular viewing. While driving, they used their own monocular bioptic telescopes to read information on highway road signs (n = 71) and pressed the horn when they saw a pedestrian hazard (n = 50). Twenty-six of the pedestrians were programed to appear, run on the road ahead of the driver for 1s within the ring scotoma and then disappear, within the period when participants were reading signs through the bioptic. The timing of the head movement to look into and out of the bioptic was determined and events were then categorized by whether or not the pedestrian hazard was present in the scene only while using the bioptic.

**Results:** When pedestrian hazards were in the scene only while subjects were using the bioptic to read a sign, detection rates were significantly higher in binocular than monocular viewing (68% vs. 40%). However, when pedestrians when subjects had a brief view of the pedestrian either beforeor after looking through the bioptic, then detection rates did not differ in binocular and monocular viewing (78% vs. 79%). By comparison, when not using the bioptic detection rates were higher (> 90%) and reaction times were shorter (without 0.95 s vs. with 1.25 s)

**Conclusions:** Our results suggest that under binocular viewing conditions the fellow eye was able to compensate for the ring scotoma to a certain extent when subjects used a monocular telescope to read road signs; however, performance was not as good as without the bioptic.

## INTRODUCTION

In order to help low-vision people to clearly see the details of distant things, bioptic telescopes were designed as small spectacle-mounted telescopes, and allowed to be driving aids^1,2^ in 45 US states,^3^ The Netherlands,^4,5^ and the province of Quebec, Canada.^6^ When people with central vision loss drive wearing bioptic telescopes, most of the time, they look below the telescope to see the traffic and road ahead. Only when the details of distant characters or traffic conditions are required, such as reading roadside signs, they tilt their heads down to look through the telescope.^7^ This kind of glance occupies less than 1.5% of the whole driving time.^8,9^

In general, there are different kinds of bioptic telescopes, such as monocular and binocular telescopes, Keplerian and Galilean telescopes.^10^ Different states have different requirements on the bioptic for driving. For example, in Massachusetts, the bioptic must be monocular, fixed focus, no greater than 3× magnification, and must be an integral part of the lens.^11^ According to the questionnaires collected from 58 bioptic drivers, almost all (95%) of them used a monocular telescope, which has only one telescope on left/right eye.^8^

When a wearer looks through a bioptic telescope, the magnified field of view leads to a ring blind area in the telescope eye, called ring scotoma, which may bring in safety risks if hazards fall in the scotoma. Therefore, the safety of driving with bioptic telescopes is a controversy over the past several years.^12^ Some have maintained that the scotoma obscures hazards ahead of the driver,^13,14^ while others have argued that the non-telescope eye, called fellow eye, in binocular viewing conditions.^15-17^ For example, in the simple visual conditions of traditional perimetry, the only blind area in the binocular view is the overlap of the physiological blind spot of the fellow eye with the ring scotoma of the telescope eye.^18^ However, in visually complex environments, such as encountered when driving, the magnification difference between the two eyes may cause binocular rivalry or suppression.^19-21^ The increase in motion velocity in the magnified view, compared with the unmagnified fellow eye view, may increase the predominance of the telescope eye^19^ which might reduce the likelihood of the fellow eye being able to compensate for the ring scotoma.

In our previous studies, when using static stimuli over a stationary patterned background, the fellow eye compensated fully for the ring scotoma,^18,22^ while for real world driving videos, only about 50% of hazards were detected by the fellow eye.^23^ Recently we developed a more realistic driving simulator paradigm to investigate hazard detection when using a bioptic telescope.^27^ The paradigm addressed some of the limitations of the earlier studies (such as prolonged viewing through the bioptic and lack of engagement in a driving task). Results of an initial study^27^ using the new paradigm suggested that the fellow eye of normally-sighted observers with simulated visual acuity loss was largely able to compensate for the ring scotoma. We now extend our investigation to visually-impaired bioptic users.

In this study, bioptic users with a wide range of driving experience were recruited representing the range of bioptic users who might be encountered in a clinical setting. In the driving simulator task some pedestrian hazards in the area of the ring scotoma were in the scene only while participants were looking through their bioptic to read information from a road sign (All-during-Tx) while others appeared before the head tilt down into the bioptic and/or remained in the scene until after the end of the upward head movement (Part-during-Tx). For All-during-Tx events we tested the hypothesis that detection rates would be higher in binocular viewing when the fellow eye could compensate than monocular viewing when the fellow eye was patched and could not compensate. For Part-during-Tx events, we expected that detection rates would not differ in binocular and monocular viewing because in both situations participants would have a brief glimpse of the pedestrian either before or after the telescope use.

## METHODS

### Participants

People with reduced VA were recruited from a database of subjects who had participated in prior studies at Schepens Eye Research Institute, from referrals from vision rehabilitation clinics, and through social media advertisements. Inclusion criteria were VA of 20/40 to 20/200 in the telescope eye (without the telescope), VA of at least 20/502 in the fellow (non-telescope) eye, current bioptic telescope user, and no manifest strabismus. In total, 16 participants were recruited, including 10 current bioptic drivers, two former bioptic drivers who stopped driving within the last three years, two former bioptic drivers who stopped driving a long time ago (17 and 25 years ago) and two bioptic users with no driving experience. Thus participants covered the entire range of bioptic drivers from those with no driving experience to those with a lot of driving experience, as might be encountered in clinical practice. A majority (88%) of the participants used their own monocular bioptic telescopes when driving in the simulator, except two who only owned binocular bioptic telescopes and used a 3x monocular bioptic telescope provided for the purposes of the study. Telescopes were focused at the distance of the driving simulator screen either using adjustable focus, when available, or lens caps.

The study followed the tenets of the Declaration of Helsinki and was approved by the institutional review board of Massachusetts Eye and Ear. Written informed consent was obtained from all participants.

#### Vision measures

Vision measures were conducted before driving tests, including visual acuity (VA), contrast sensitivity, and visual fields. Using Test Chart 2000 Pro (Thomson Software Solutions, Herts, England), VA was measured monocularly and binocularly without the bioptic and with the telescope eye through bioptic, at a distance of 20 feet. In addition, binocular letter contrast sensitivity without bioptic was measured using the Mars Chart (The Mars Perceptrix Corporation, Chappaqua, New York).

A custom computerized test^24^ was used to carry out tangent-screen kinetic perimetry to map the area of the ring scotoma with white stimuli presented against a black background. The participant, sitting at 1 meter from a rear-projection screen (1.65 × 1.25 m^2^), was asked to focus on a cross (0.2° × 1.2°) in the center through the bioptic telescope, and then to press a button as soon as a square stimulus (1.9°) was seen. During the test, the square was moved inwards or outwards from unseen to seen in order to plot the inner or outer boundary of the ring scotoma. For all the participants the monocular ring scotoma was measured with the fellow eye patched (Figure SS). In addition, the binocular visual field with the telescope eye viewing through the bioptic was also measured when the subject had central visual field loss in the fellow eye that might overlap with the ring scotoma in the telescope eye to create a scotoma in the binocular visual field (Figure SD), or when the telescope might cause a scotoma in the binocular visual field (such as the Ocutech VES Keplerian telescope; Figure ST), or.

**Figure SS.**
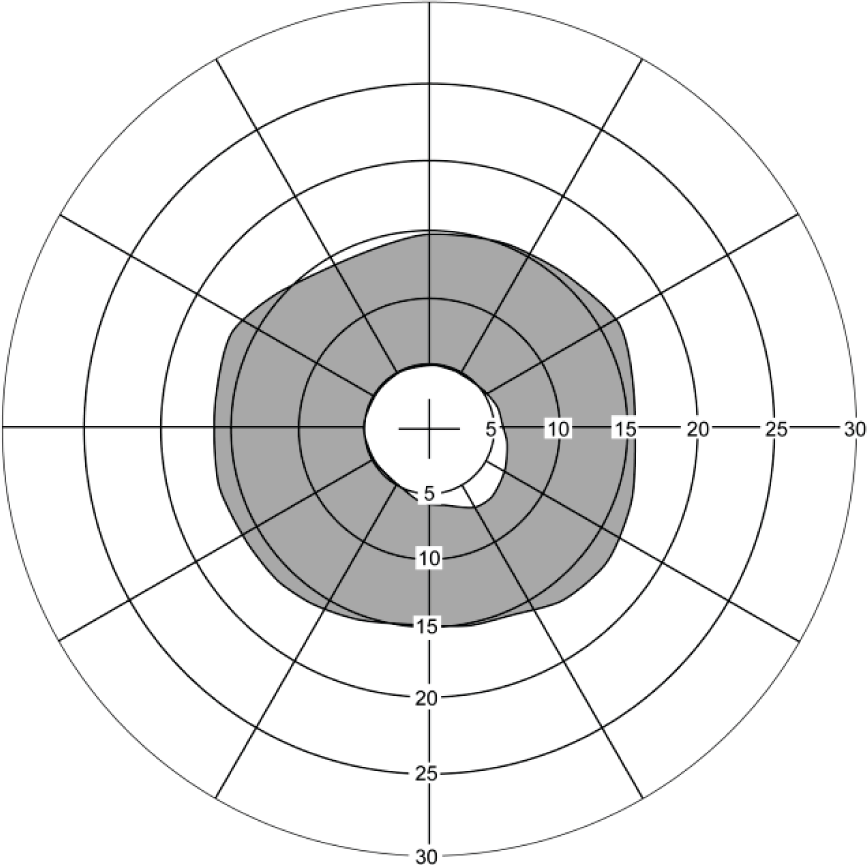
Ring scotoma area when looking into 3.0× monocular bioptic telescope. The central white area was the magnified field of view, which the participant saw through telescope, while the outer white area was the field of view outside the telescope (without magnification). The grey shading area was the ring scotoma, usually located at around 5° to 15° when looking through 3.0× monocular bioptic telescope.

**Figure SD.**
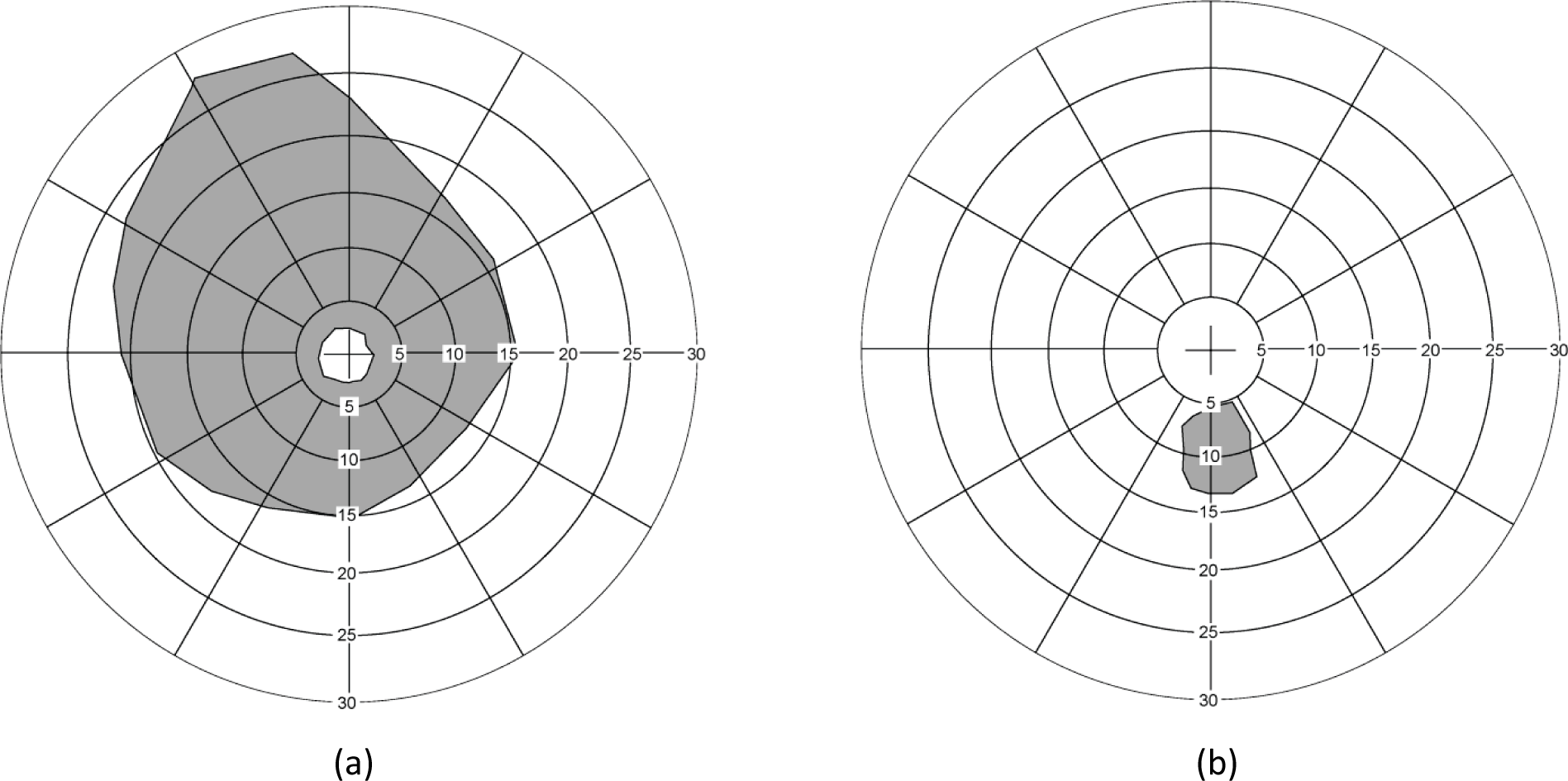
Monocular and binocular visual fields for a participant with macular holes looking through a 3.0× Galilean monocular telescope. (a) Ring scotoma in the monocular visual field of the telescope eye (fellow eye patched); (b) Scotoma in the inferior binocular visual field where the ring scotoma of the telescope eye overlapped the scotoma from the macula hole in the fellow eye.

**Figure ST.**
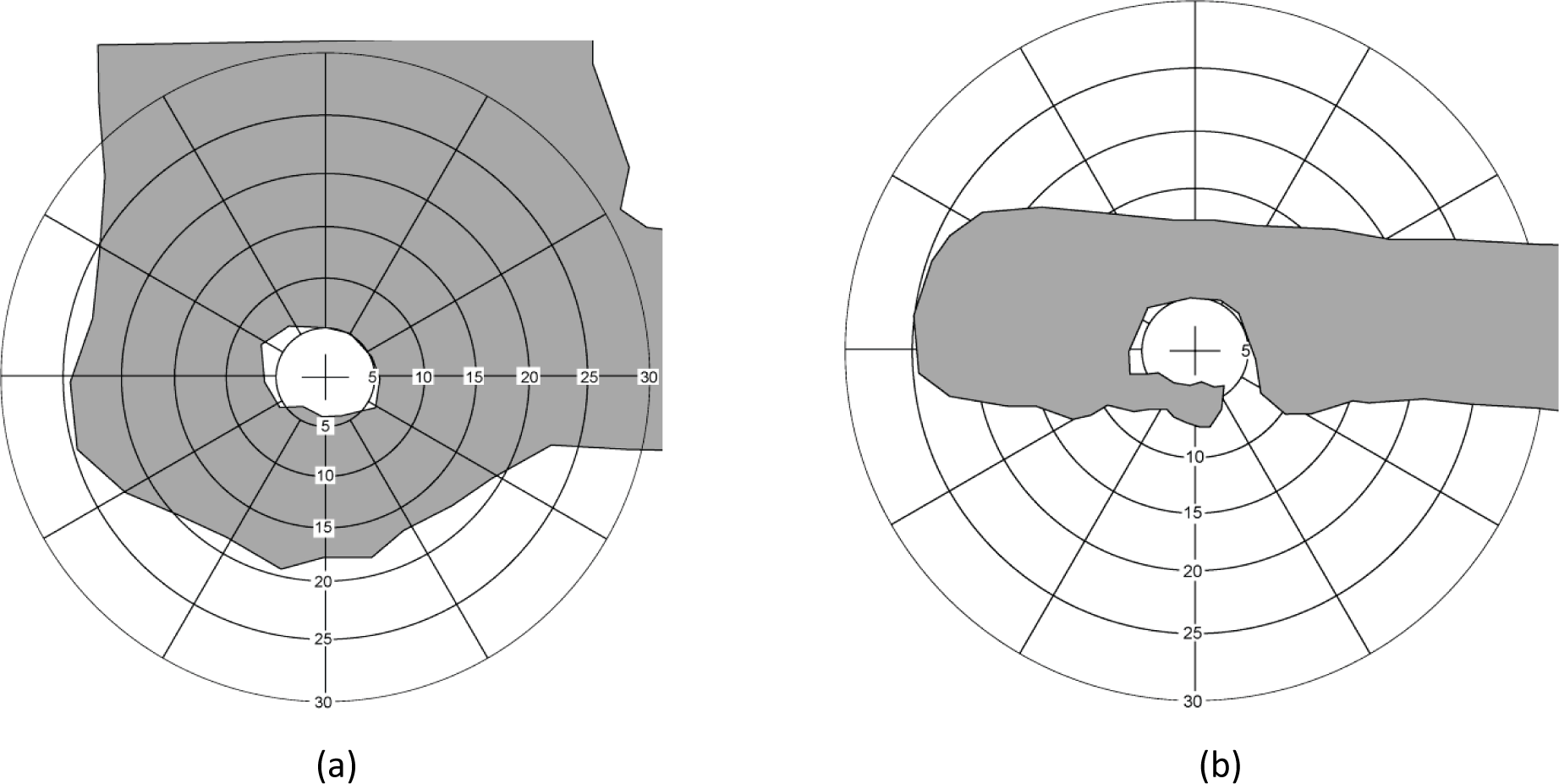
Monocular and binocular visual fields for a participant wearing Ocutech VES 3x Keplerian bioptic telescope in which the rectangular housing of the telescope is mounted across the top of both spectacle lenses. (a) Ring scotoma in the monocular visual field of the telescope eye (fellow eye patched); (b) Large scotoma in the binocular visual field caused by the rectangular housing of the telescope.

#### Driving simulator task

##### Apparatus

The driving simulator was a DE-1500 (FAAC Corp., Ann Arbor, MI), with five LCD monitors (42” diagonal, 1366 × 768 pixels, 60Hz) providing 225° field of view. The simulator included all the controls found in an automatic transmission vehicle, and a 3 degrees-of-freedom motion seat. Driving scenarios were developed using the Scenario Toolbox software (FAAC Corp., Ann Arbor, MI). The location and status of the driver’s car in the virtual world, as well as the data of all programmed objects, such as the pedestrians, the road signs and other cars, were continuously recorded at 30Hz.

A Smart Eye^®^ remote six-camera IR system (Smart Eye Pro 6.1, Gothenburg, Sweden) was used to track the participant’s head and eye movements at 60Hz. The timing of the bioptic telescope use was determined from the head movement data based on tracking of facial features which could be tracked reliably even when using the bioptic. Although eye movements were recorded, the data were often very noisy with a lot of data drop outs due to difficulties in tracking eyes of people with nystagmus, tracking through high prescription glasses and loss of tracking when looking into the bioptic. Thus the eye data were not used in analyses. Custom software was used to synchronize the 60-Hz Smart Eye data stream with the 30-Hz simulator data stream and the virtual world coordinate system.

##### Driving simulator procedure

Before experimental data collection, each subject took at least 30 minutes’ driving practice in the driving simulator, in order to become familiar with vehicle control, reading road signs through bioptic telescope in the simulator and driving with the fellow eye patched. Once participants were acclimated to the simulator, they completed 6 test drives, each about 10 minutes, on rural roads with light oncoming traffic. The six test drives included the three used in the prior study^27^ with simulated vision impairment and an additional three developed for the current study using the same criteria. Three drives were undertaken with binocular viewing (fellow eye was open) and three drives with monocular viewing (fellow eye was patched). Binocular and monocular drives were interleaved with the order counterbalanced across subjects. To ensure consistency of event timing, a speed cap was set at 35 mph and participants were asked to drive as close as possible to this maximum speed. Participants could increase speed up to the maximum using the accelerator and could slow down the car with brakes when necessary. They also had full control of vehicle steering. Participants were asked to use their bioptic to read information from road signs, to press the horn on the steering wheel as soon as they detected any pedestrian, either running or stationary, and to obey all the normal rules of the road. For all participants, a 5-point calibration of the SmartEye^®^ tracker was performed before experimental data collection commenced.

Former bioptic drivers and participants with no driving experience were given additional time, as needed, to practice driving in the simulator. The two participants without driving experience learned to control the speed and steering of the virtual vehicle very quickly as it was not dissimilar to a video game. The driving scenarios did not involve making any turns at intersections or interactions with other vehicles; therefore these driving skills were not included in the driving training provided to the two non-drivers. Experimental data collection did not commence until the participant demonstrated sufficiently good control of vehicle speed and steering.

##### Sign-reading and hazard-detection tasks

In the six test drives, besides vehicle control, there were two main tasks for low-vision participants: sign reading and pedestrian detection. Details of the tasks are given elsewhere.^27^ In brief, directional road signs were designed according to Standard Highway Signs in color, font, and text spacing (refer to the Manual of Uniform Traffic Control Devices http://mutcd.fhwa.dot.gov/ser-shs_millennium.htm). The custom navigation information on a road sign consisted of the road name (“Massachusetts Pike” or “Massachusetts Ave”) and the distance (“0.8 Miles” or “0.3 Miles”). The signs appeared at pseudo-random time intervals when the participant’s car was about 88 m away from the programmed sign locations. When a highway sign appeared, the participant was asked to identify the words on it through the bioptic telescope, and verbally report “Pike” or “Ave” and “8” or “3”.

In addition to the road signs, pedestrians were programmed to appear at pseudo-random time intervals, including pedestrians standing at the side of the road and pedestrians running across the participant’s driving lane from left to right or from right to left. Across the 6 drives there were a total of 26 occasions when a running pedestrian appeared at the same time as a highway sign (*sign+pedestrian* events). The pedestrian was programmed to appear approximately 1 second after the highway sign when the participant was expected to have already dipped their head to look into the bioptic.^27^ The pedestrian appeared in an area of the lower visual field that would be within the ring scotoma when reading the sign through the bioptic (Figure SP). It ran for about 1 s ahead of the participant’s car and then disappeared before the sign-reading task was anticipated to be finished. Whether the pedestrian was totally within the area of the ring scotoma on any particular trial depended on vehicle heading and speed, as well as when and for how long participants looked through the telescope.

**Figure SP:**
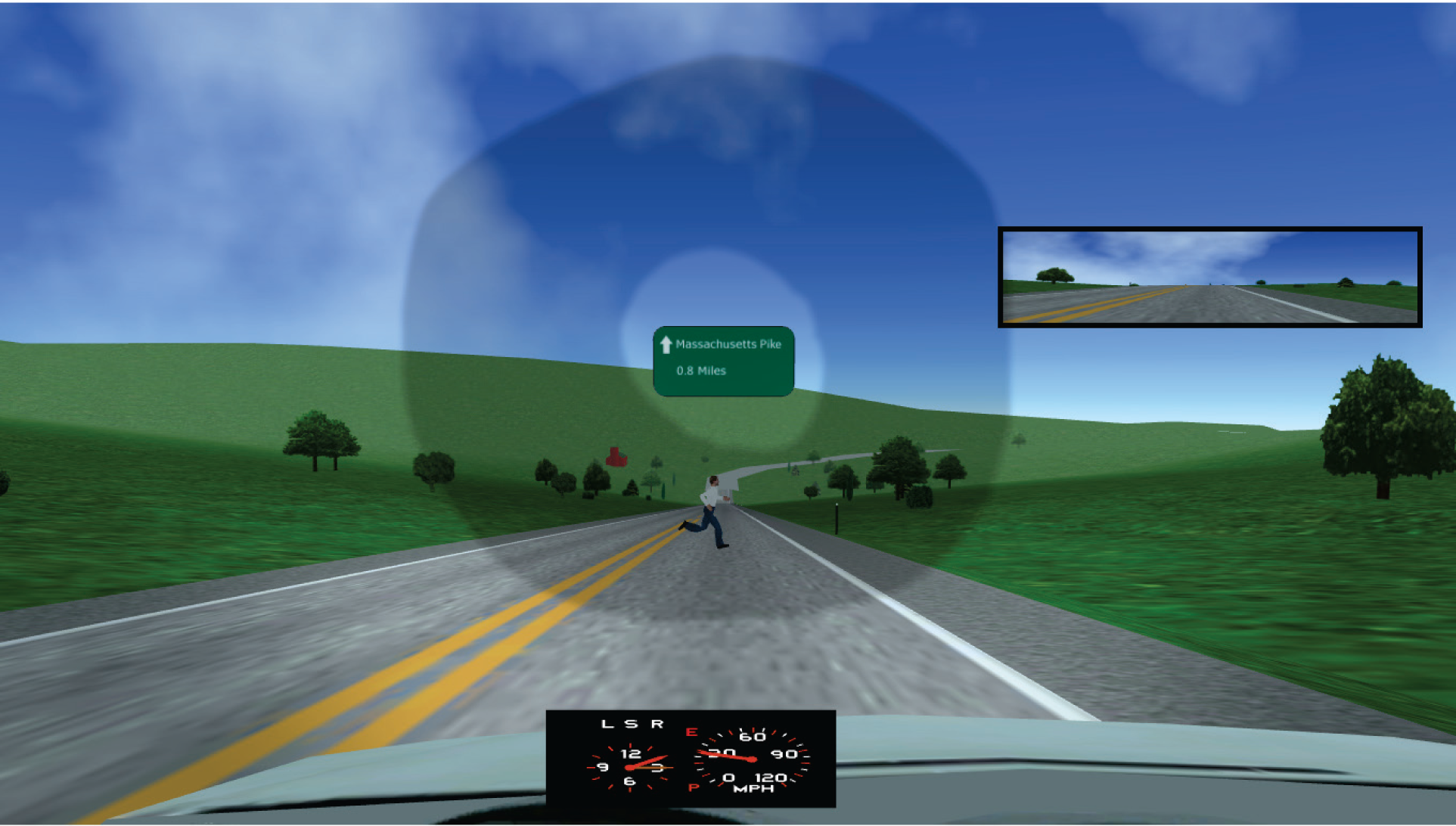
Screen-shot of a *sign+pedestrian* event, in which the vehicle was about 70 m from the sign and 35 m from the pedestrian. The pedestrian was programmed to appear, run across the road ahead of the driver for about 1 s, and disappear, all while the participant was reading the sign through the bioptic. The pedestrian was within the ring scotoma area in the monocular visual field (shown by grey shading). The small area around the sign, without shading, is the magnified field of view through telescope. Only the central monitor of the simulator is shown; the inset on upper right is the rearview mirror.

To keep the participants naïve about the purpose of this study, a variety of events were programmed. In addition to the 26 *sign+pedestrian*events in the six test drives, there were 35 signs without a pedestrian, 14 running pedestrians without a sign, and 33 stationary pedestrians without a sign. Only responses to the *sign+pedestrian*events and running pedestrians without a sign were analyzed.

##### Timing of sign+pedestrian events

For each *sign+pedestrian* event, a plot of vertical head position with the timings of sign appearance, pedestrian appearance and disappearance was generated (Figure HT). The timing of the bioptic use was then determined from the vertical head position data using a custom MATLAB program based on data cursor mode. The program was used to mark and save the coordinates of the time points at which the downward head movement started and the upward head movement finished, T1 and T2, respectively on Figure HT.

**Figure HT.**
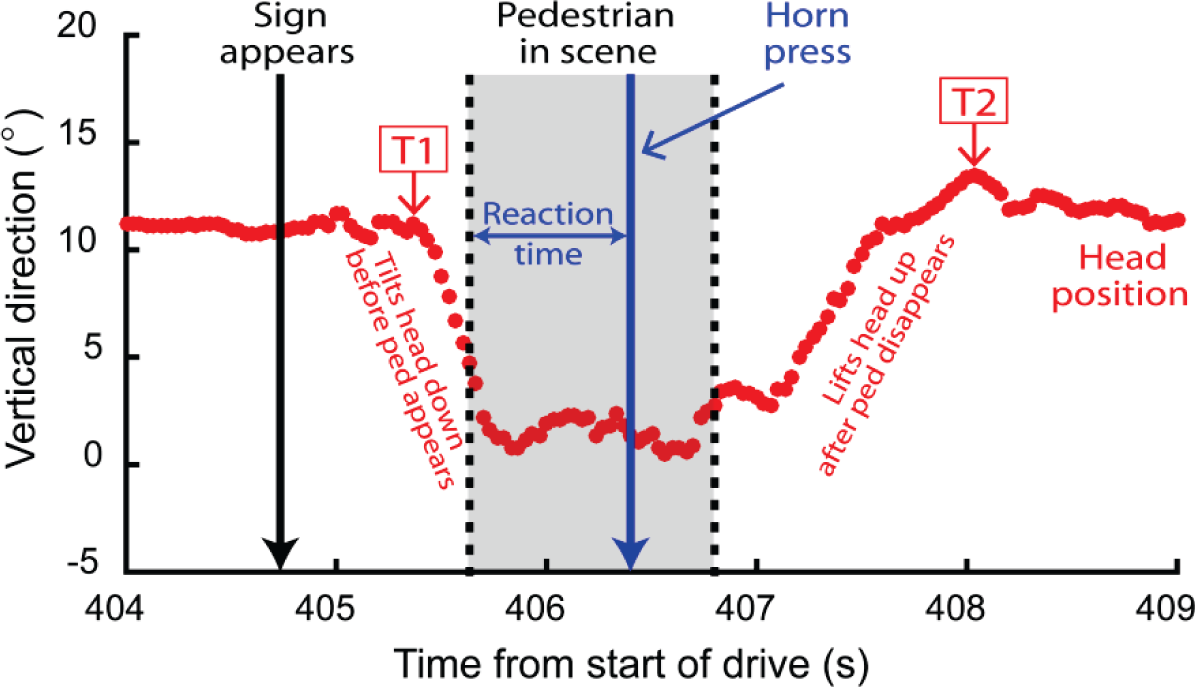
Vertical head position and event timing for an “All-during-Tx” sign+pedestrian event in the binocular viewing condition. The participant started to tilt their head down to look through the bioptic about 0.6 s after the sign appeared with the end of the upward head movement about 2.5 s later. Since the pedestrian was in the scene (grey-shaded region) only while the subject was looking into telescope, this event was classified as an All-during-Tx event. In this example, the pedestrian was detected (as indicated by the horn press < 1 s after pedestrian appearance), because in binocular viewing, the fellow eye was open and able to compensate for the ring scotoma. T1 marks the start of the downward tilt. T2 marks the end of the upwards movement.

For further analysis of sign+pedestrian events, five timing variables were calculated, including (1) the time from sign appearance to pedestrian appearance, (2) the duration of pedestrian in scene, (3) the time from sign appearance to T1, the start of the head tilt down, (4) the time from T1 to the pedestrian appearance, and (5) the time from pedestrian disappearance to T2, the end of the upward head movement. The first two variables were used to verify the consistency of the pedestrian timing with respect to the sign appearance across events; the third characterized the timing of bioptic use; and the last two were used to classify sign+pedestrian events.

##### Categorizing sign+pedestrian events

Sign+pedestrian events were categorized with respect to the amount of overlap between the period of bioptic telescope use and the period of time for which the pedestrian was in the scene. Events were categorised into 4 types: 1) No head dip – the participant did not look into the telescope; 2) No overlap – the participant looked into the telescope, but the period of telescope use did not overlap with the time that the pedestrian was in the scene; 3) Part-during-Tx – there was some overlap between the telescope use and the pedestrian but the pedestrian either appeared before the start of the telescope use or disappeared after the end of the telescope use (Figure PT); and 4) All-during-Tx – the driver was engaged in the bioptic telescope through the entire period of time the pedestrian was in the scene i.e., the pedestrian appeared after the start and before the end of the telescope use (Figure HT).

**Figure PT:**
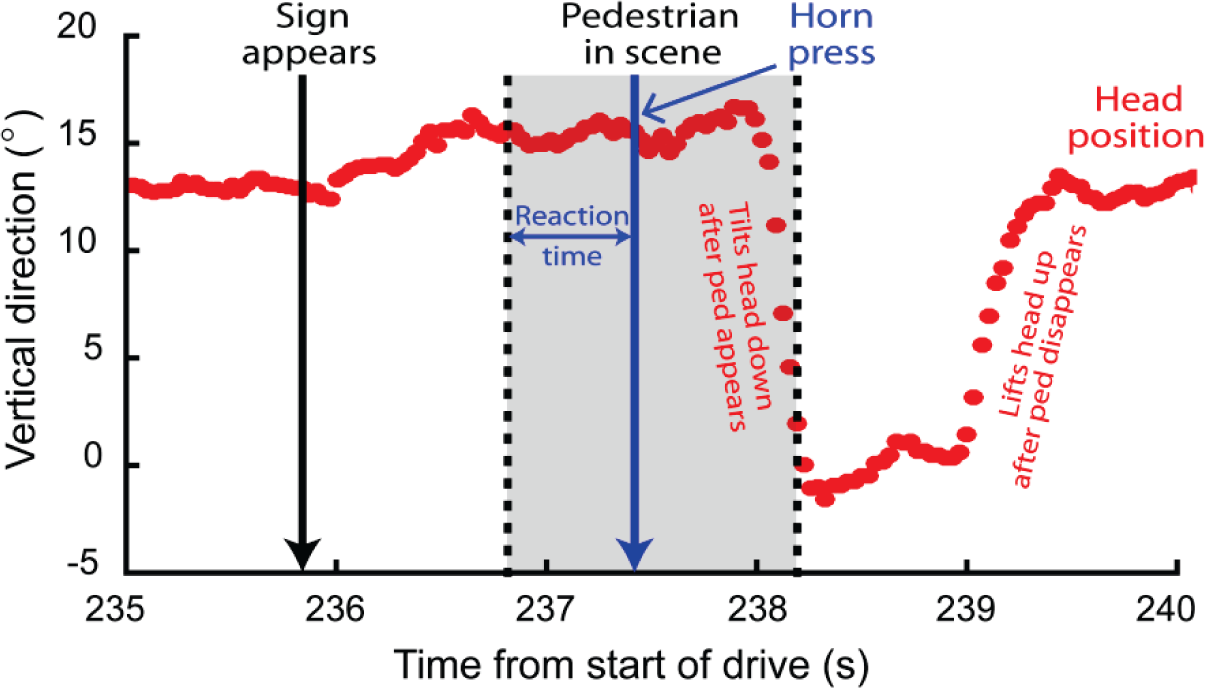
Vertical head position and event timing for a “Part-during-Tx” sign+pedestrian event in the monocular viewing condition. The pedestrian appeared about 1 s after the sign. However, the participant did not start the downward head movement until 1.1 s later such that there was only a brief period of overlap (0.2 s) between the telescope use and the pedestrian being in the scene. Thus, the event was categorized as Part-during-Tx. In this case, the view of the pedestrian before looking into the telescope was sufficient for detection by the telescope eye despite the monocular viewing conditions.

The period of telescope use was defined as being the time between the eyes beginning to move up into the telescope and completing their downward movement out of the telescope. In the prior study we were unable to track the eyes through the diffusing filters used to create the simulated vision impairment and therefore could not base our analysis of the start and end of telescope use on the eye movements. Instead we used the start and end of the head movement (T1 and T2, respectively). For the majority of subjects with vision impairment in the current study, eye movements were also not tracked sufficiently well to be able to define the start and end of the telescope use. We therefore recorded eye and head data from a group of normally-sighted participants who used a monocular bioptic to read the highway signs while driving in the simulator (but without the diffusing filters on the carrier lens). From these recordings, we established that the eye started to move up about xx ms before the start of the downward head movement (T1) while at the end of the telescope use, the eye completed the downward movement on average about yy ms before the end of the upward head movement (T2). For the categorization of the sign+pedestrian events we therefore defined the start of the telescope use as xx ms before T1 and the end of the telescope use as yy ms before T2.

### Statistical analysis

Head movement data were available for 413 of the total 416 sign+pedestrian events. A mixed effects binary logistic regression analysis was used to evaluate the effects of viewing condition (monocular or binocular), event category (All-during-Tx or Part-during-Tx) and their interaction on whether pedestrians were detected in the sign+pedestrian events. Driving status (current or non-current) and binocular field status (scotoma present/absent in the inferior field) were also included as fixed factors because both these variables might affect detection of pedestrian hazards. Subject was included as a random factor.

Reaction time data for events where the pedestrian was detected did not conform to a normal distribution, even after a log transform. Median reaction times were therefore computed for each subject for All-during-Tx, Part-during-Tx and no-sign pedestrian events. The medians did not differ from a normal distribution and were therefore analyzed with parametric statistics (repeated measures ANOVA and paired t-tests).

The five timing variables which were used to characterize bioptic use were analyzed for sign+pedestrian events. Median values were computed for each subject for each variable and the medians were analyzed with non-parametric statistics. All statistical analyses were performed with STATA/IC 14 (College Station, TX). A value of α = 0.05 was taken to indicate statistical significance.

## RESULTS

### Sample characteristics

The 16 participants (50 % male) ranged in age from 17 to 80 years (median 52.5 years). The majority had congenital vision loss including albinism (n=6), congenital cataracts (n = 3), nystagmus (n = 2) and retinopathy of prematurity (n = 1). The four participants with adult onset vision loss had optic nerve damage (n = 1), macular holes (n = 1), Stargardts (n = 1) and age-related macular degeneration (n = 1). For the telescope eye, median visual acuity was 20/100 without the bioptic and 20/36 with the bioptic. Median acuity of the fellow eye was 20/126. Four participants had a scotoma in the inferior binocular field when viewing through the telescope. For one subject the scotoma was a result of macular disease (Figure SD) while for the other three subjects, the scotoma resulted from the rectangular housing of a Ocutech VES bioptic telescope (Figure ST). Three of the four participants who had stopped driving had a scotoma in the lower binocular field (one had a disease scotoma and two had device scotomas).

The majority of participants used their bioptic telescope every day (10/16) and reported that it was very helpful (12/16). It was median 20 years (range 1 to 46 years) since their first bioptic telescope had been prescribed. The majority (12/16) of participants used Galilean telescopes, mostly manufactured by Ocutech (8/16) with 3.0× the most common level of magnification (range 2.2× to 6.0×).

Of the 10 current drivers, 7 drove more than 100 miles per week (median, IQR). All the drivers thought the bioptic telescope was helpful in driving, with 80% reporting it was very helpful. The main tasks for which the bioptic was used included: reading road/traffic signs (90%), seeing traffic light signals (90%), reading street name signs (80%), seeing pedestrians crossing the road / hazards ahead (80%), and judging when it was safe to turn at an intersection without traffic lights (80%). Less common tasks included seeing brake lights and signal lights on cars in front (40%), judging the distance to the car in front (40%), and judging when it was safe to overtake another car (30%).

### Sign reading performance

In the sign reading task, the correct response rate was the proportion of signs, on which both the street name and the distance were identified correctly. All the participants had relatively high correct response rates (ranged from 80%-100% across participants). Additionally, the correct response rates were not different in monocular and binocular viewing conditions (median 94% and 97% respectively, p = 0.06), and for the tasks with vs. without pedestrians (median 97% and 96% respectively, p = 0.209).

### No-sign events detection performance

Detection rates for pedestrians not coincident with signs (no-sign events) were high in both the monocular and binocular viewing conditions (94% and 97%, respectively, Figure DR) and reaction times did not differ between the two conditions (medians 0.975 s and 0.9 s, respectively).

### Sign+pedestrian events

#### Bioptic use timing and event categorization

In all the sign+pedestrian events, median time from sign appearance to pedestrian appearance was 1 s (IQR 0.9-1.0 s), and median time from pedestrian appearance to pedestrian disappearance was 1.1 s (IQR 1.0-1.2 s). Thus, the timing of sign+pedestrian events was very consistent across subjects.

Regarding bioptic use, the median time from sign appearance to the start of the head tilt down was 1.4 s with the majority of head tilts starting between 1.0 and 1.8 s after the sign appearance (Figure TS). The majority (77%) of pedestrians appeared after the head tilt down when the participants were already looking through bioptic telescopes (Figure TP a). And 72% pedestrians disappeared before the end of the head lift up while the participants were still looking into the bioptic telescope (Figure TP b).

**Figure TS.**
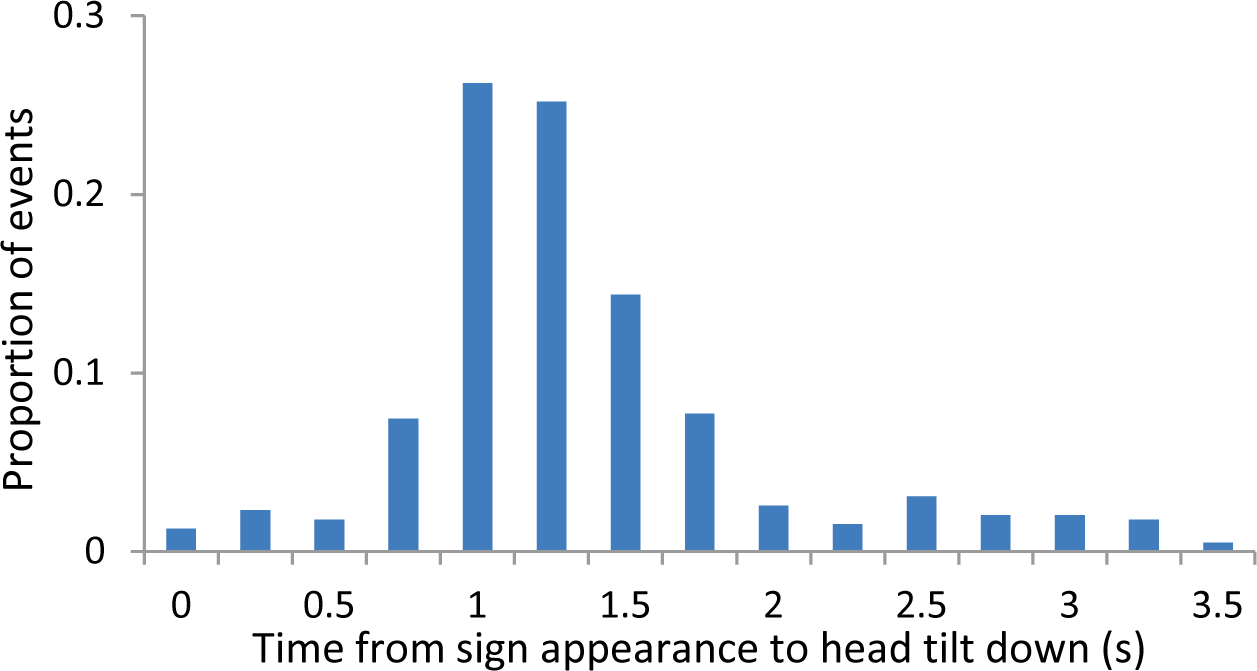
Distribution of time from sign appearance to head tilt down.

**Figure TP.**
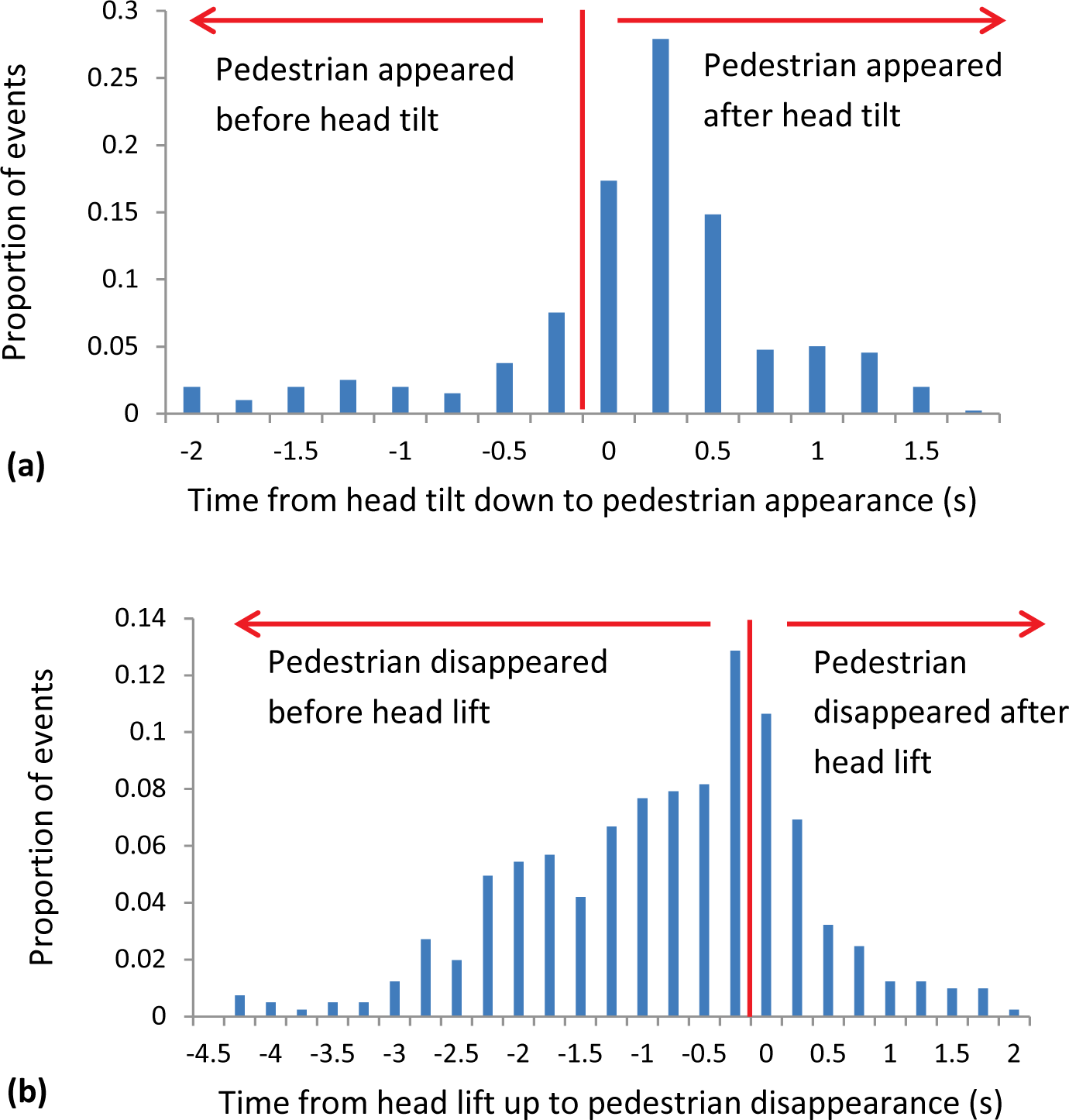
Timing of pedestrian and bioptic use. (a) Distribution of time from head tilt down to pedestrian appearance; (b) distribution of time from pedestrain disappearance to head lift up.

Across all 16 participants, there were 413 *sign+pedestrian* events including 35 (8%) with no head tilt (telescope was not used), 44 (11%) with no overlap between the period of bioptic use and the time when the pedestrian was in the scene (no overlap), 147 (36%) when the pedestrian partly overlapped with the telescope use (Part-during-Tx) and 187 (45%) when the pedestrian was only in the scene while the participant was looking through the telescope (All-during-Tx). The proportions of each kind of event did not differ between monocular and binocular viewing (Figure PE). In All-during-Tx events, pedestrians appeared median 0.4 s after head tilt down, while in Part-during-Tx events, pedestrians appeared median 0.27 s before head tilt down.

**Figure PE.**
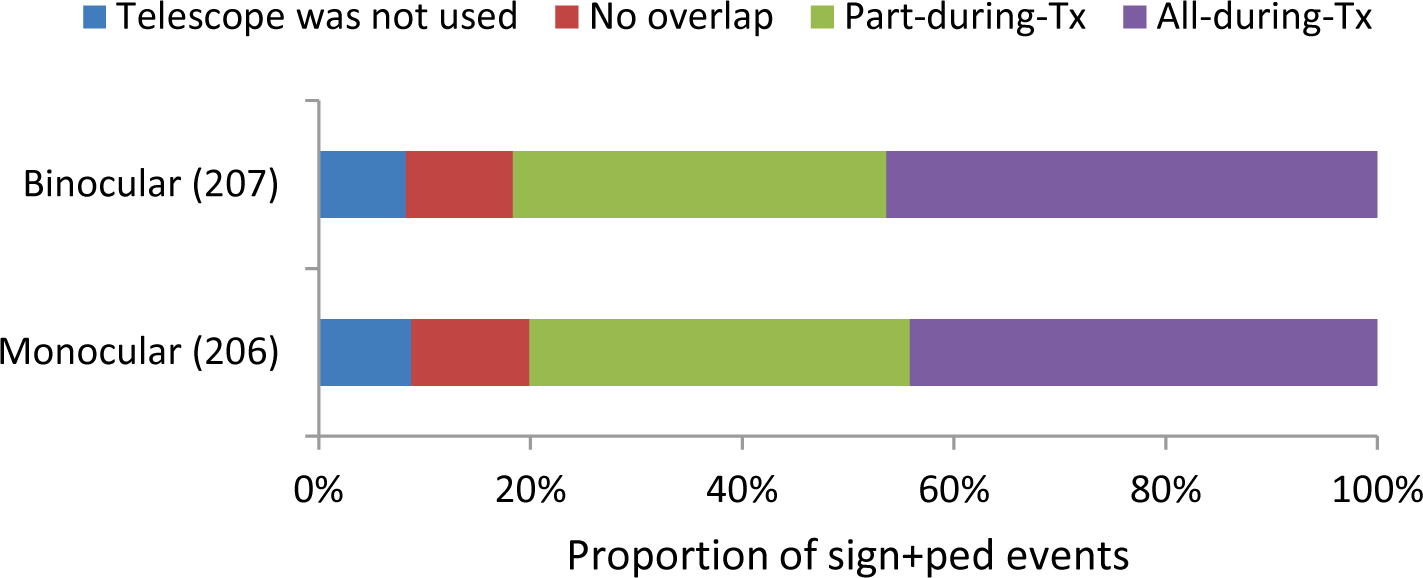
Proportion of sign+pedestrian events that were categorized as Telescope was not used, No overlap, Part-during-Tx and All-during-Tx in monocular and binocular viewing conditions. There were totally 206 and 207 sign+pedestrian events in monocular and binocular viewing, respectively.

#### Pedestrian detection rates

For All-during-Tx events, mean detection rates were significantly higher in binocular than monocular viewing (68% vs. 40%; Figure DR). However, for Part-during-Tx events, mean detection rates did not differ in binocular and monocular viewing (78% vs. 79%). By comparison, detection rates for no-overlap events were higher and did not differ from detection rates for no-sign events (Figure DR).

**Figure DR.**
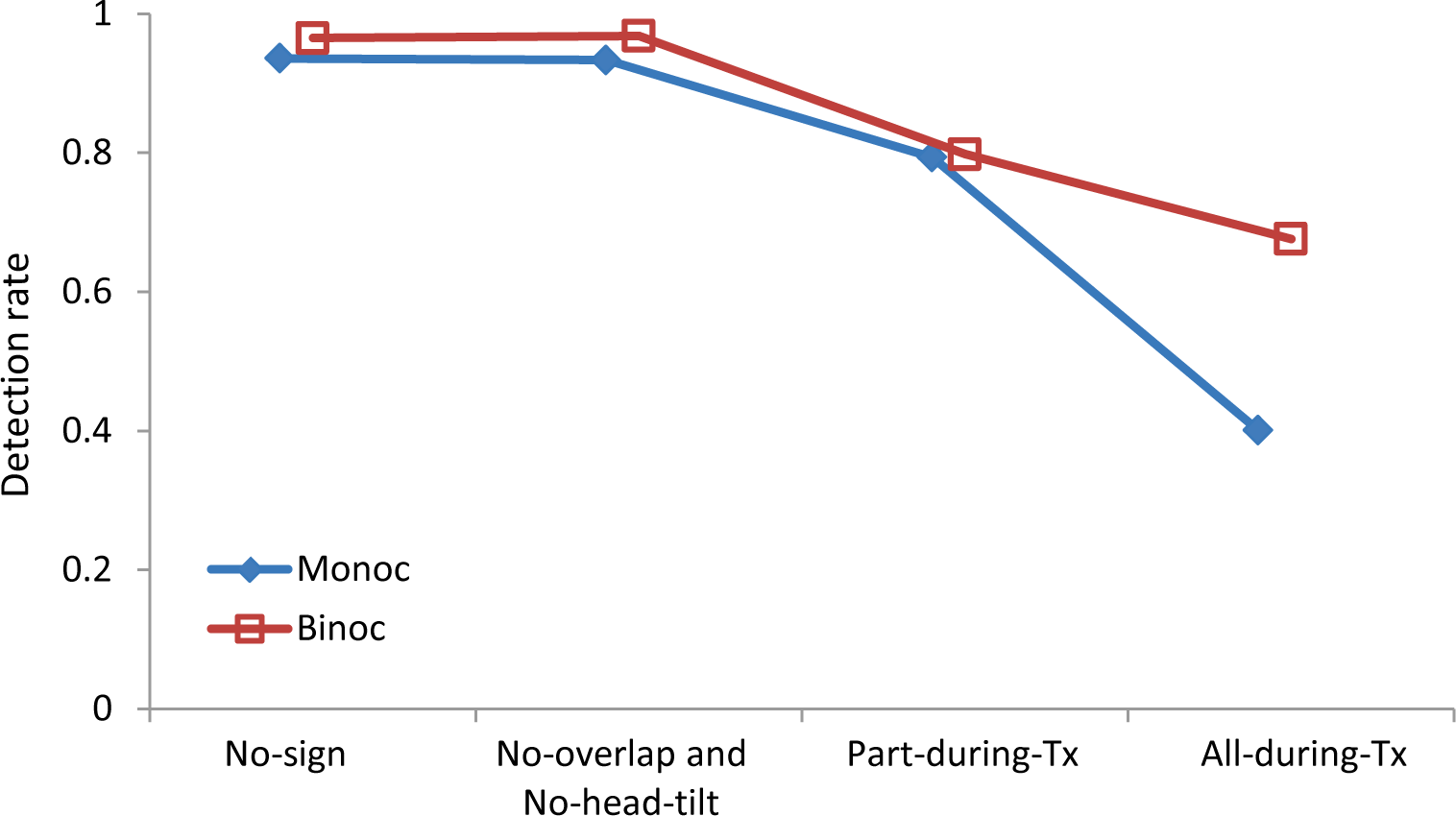
Mean detection rates for No-sign, No-overlap and no-head-tilt, Part-during-Tx and All-during-Tx events in binocular and monocular viewing conditions.

#### Reaction times

Figure RT shows the reaction times to pedestrians. Reaction times did not differ in binocular and monocular viewing conditions and were therefore pooled in analyses. Compared to the No-sign pedestrian events, reaction times were significantly longer for Part-during-Tx and All-during-Tx sign+pedestrian events (p < 0.001), but did not differ from reaction times in No-overlap events (p = 0.82). There was a trend for reaction times to be longer for All-during-Tx than Part-during-Tx events (p = 0.06).

**Figure RT.**
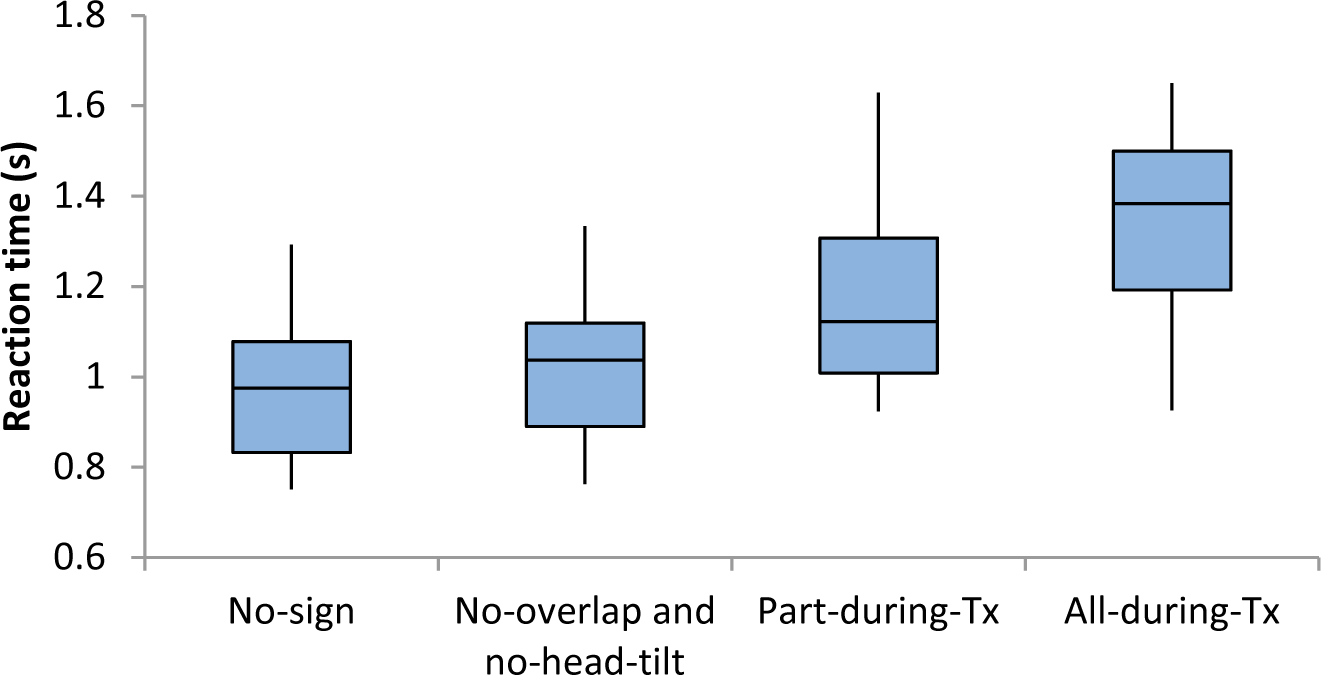
Boxplots of reaction times for No-sign, No-overlap and no-head-tilt, Part-during-Tx and All-during-Tx events. Horizontal line within the box is the median; the vertical extent of the box is the IQR; whiskers represent the data range with outlier excluded.

### Effects of driving status

The effects of driving status (current, former, non-driver) were evaluated for bioptic use timing variables and detection performance. We expected that current drivers would have more efficient bioptic use behaviors when reading road signs and that they might show increased ability of the fellow eye to compensate in binocular viewing conditions.

Inspection of the bioptic use timing data suggested that non-drivers and former drivers behaved in a similar manner and were therefore combined into one group for analysis (non-current driver).

There was no effect of driving status (current vs. non-current) on the time between the sign appearing and the start of the head tilt down (Table DSA; p = 0.83). However, current drivers looked through the telescope for a significantly shorter period of time than non-current drivers (p = 0.015) and also took less time to complete both the downward (p = 0.007) and upward (p = 0.10) head movements (Table DSA).

**Table DSA:** Median times for bioptic use timing variables for non-current and current drivers

| Driving status | Time from sign to head tilt down (s) | Total duration of bioptic use (s) | Time for downward head movement (s) | Time for upward head movement (s) |
| --- | --- | --- | --- | --- |
| Non-current | 0.75 | 2.95 | 0.50 | 0.70 |
| Current | 0.82 | 1.66 | 0.38 | 0.48 |

For detection rates, all participants, irrespective of whether they were current, former or non-drivers detected the majority (> 90%) of no-sign pedestrians Table DSB). For sign+pedestiran events, the effects of driving status were only analyzed for the All-during-Tx events because there were insufficient numbers of Part-during-Tx events. Current drivers, recent former drivers (≤3 years) and non-drivers all showed an improvement in detection performance between monocular and binocular viewing, suggesting that even non-drivers can use the fellow eye to compensate for the ring scotoma.

**Table DSB:**
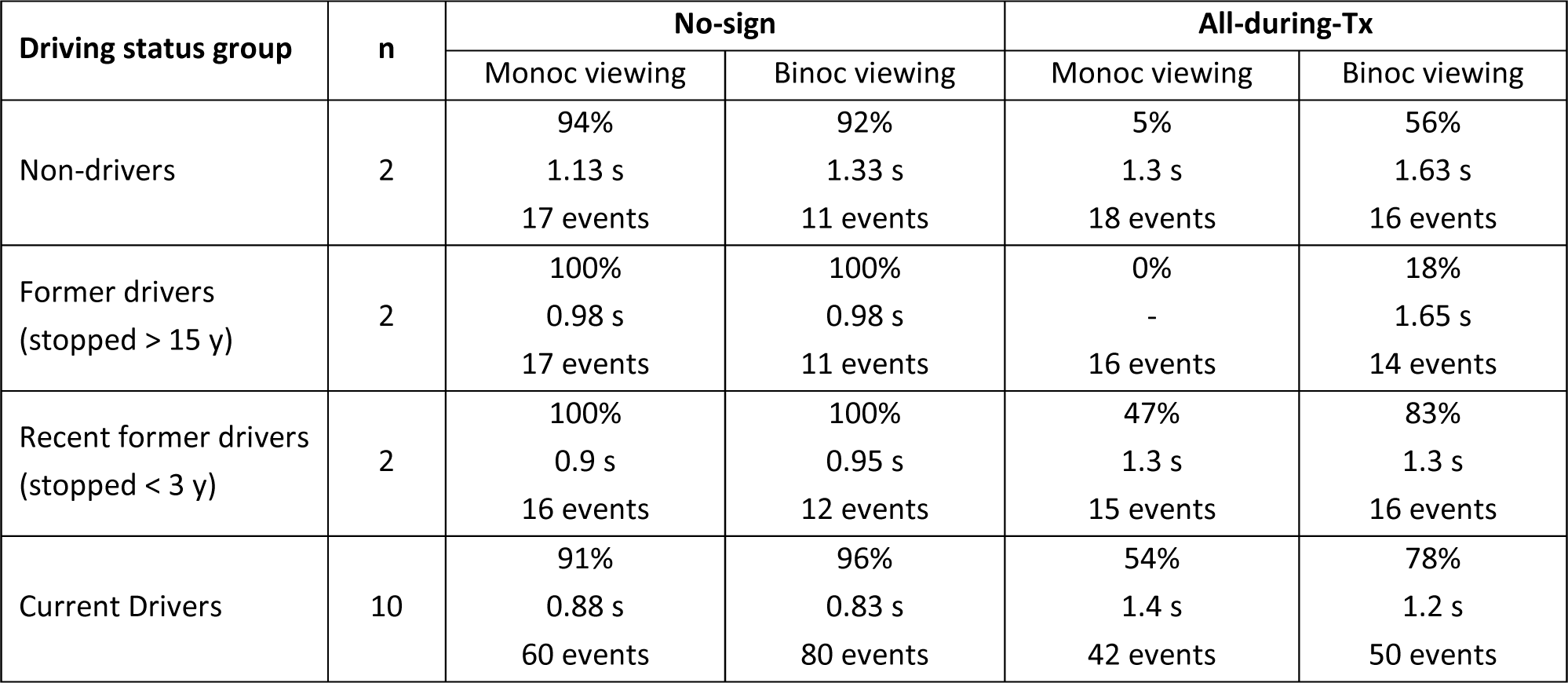
Detection rates and reaction times by driving status

| Driving status group | n | No-sign |  | All-during-Tx |  |
| --- | --- | --- | --- | --- | --- |
|  |  | Monoc viewing | Binoc viewing | Monoc viewing | Binoc viewing |
| Non-drivers | 2 | 94%<br>1.13 s<br>17 events | 92%<br>1.33 s<br>11 events | 5%<br>1.3 s<br>18 events | 56%<br>1.63 s<br>16 events |
| Former drivers<br>(stopped > 15 y) | 2 | 100%<br>0.98 s<br>17 events | 100%<br>0.98 s<br>11 events | 0%<br>-<br>16 events | 18%<br>1.65 s<br>14 events |
| Recent former drivers<br>(stopped < 3 y) | 2 | 100%<br>0.9 s<br>16 events | 100%<br>0.95 s<br>12 events | 47%<br>1.3 s<br>15 events | 83%<br>1.3 s<br>16 events |
| Current Drivers | 10 | 91%<br>0.88 s<br>60 events | 96%<br>0.83 s<br>80 events | 54%<br>1.4 s<br>42 events | 78%<br>1.2 s<br>50 events |

Interestingly, however, non-drivers had lower detection rates than recent-former and current drivers in both monocular and binocular viewing. Former drivers who stopped driving a long time ago (> 15 years) were the only participants to show little improvement in detection rates between monocular and binocular viewing. In fact, detection rates were very low in both monocular and binocular viewing. This finding is explained by the fact that both of these participants had scotomas in the lower binocular visual field in an area that could impact detection performance.

## DISCUSSION

For All-during-Tx events, pedestrian detection rate was higher in binocular than monocular viewing (as shown in Figure DR), because the fellow eye helped to detect some pedestrians appearing in the ring scotoma of the telescope eye. For Part-during-Tx events, detection rates were not different in binocular and monocular viewing. This is because during a period of time, the pedestrian appeared outside the ring scotoma of the telescope, and thus the telescope eye could detect it in monocular viewing. These results were consistent with our hypothesis, and suggested that the fellow eye was able to compensate for the ring scotoma when driving with monocular bioptic telescopes.

Even in binocular viewing, detection rates for Part-during-Tx and All-during-Tx events were lower than for No-sign and No-overlap events. Similar results were found in our prior study in which normally-sighted participants with simulated vision impairment used a monocular bioptic telescope to perform the same sign-reading task. Taken together, these findings suggest that the additional processing resources devoted to engagement in using the telescope and/or reading the sign reduced those available for other tasks such as detection of pedestrians. Interestingly, a similar effect was found when people with impaired vision completed a reading task without a bioptic telescope while performing a video-based hazard perception task; hazard detection rates were reduced when performing the reading task. In sum, the ring scotoma appears to have little impact on pedestrian hazard detection in binocular viewing when the fellow eye can compensate. However the divided attention conditions of using the telescope and/or reading a sign coincident with a hazard appearance decreases the likelihood that the hazard will be seen. In monocular viewing conditions provided the bioptic is used only briefly and drivers have a glimpse of the hazard either before or after the telescope use (Part-during-Tx events), then hazard detection rates do not differ from those in binocular viewing.

## ACKNOWLEDGEMENTS

The authors would like to thank Robert Goldstein for programming software. Funded in part by NIH grants R01-AG04197, R01-EY025677, 1S10RR028122 and P30EY003790.

